# Using imaging photoplethysmography for heart rate estimation in non-human primates

**DOI:** 10.1101/252403

**Authors:** Anton M. Unakafov, Sebastian Möller, Igor Kagan, Alexander Gail, Stefan Treue, Fred Wolf

## Abstract

For humans and for non-human primates heart rate is a reliable indicator of an individual’s current physiological state, with applications ranging from health checks to experimental studies of cognitive and emotional state. In humans, changes in the optical properties of the skin tissue correlated with cardiac cycles (imaging photoplethysmogram, iPPG) allow non-contact estimation of heart rate by its proxy, pulse rate. Yet, there is no established simple and non-invasive technique for pulse rate measurements in awake and behaving animals. Using iPPG, we here demonstrate that pulse rate in rhesus monkeys can be accurately estimated from facial videos. We computed iPPGs from seven color facial videos of three awake head-stabilized rhesus monkeys. Pulse rate estimated from iPPGs was in good agreement with reference data from a pulse-oximeter with error of pulse rate estimation below 5% for 82% of all epochs, and below 10% for 98% of the epochs. We conclude that iPPG allows non-invasive and non-contact estimation of pulse rate in non-human primates, which is useful for physiological studies and can be used toward welfare-assessment of non-human primates in research.

## 1 Introduction

Heart rate is an important indicator of the functional status and psycho-emotional state for humans, non-human primates (NHP) [1–3] and other animals [4, 5]. A common and the most direct tool for measuring heart rate is the electrocardiogram (ECG) [6]. ECG acquisition requires application of several cutaneous electrodes, which is not always possible even for human subjects [7]. ECG acquisition in NHP is complicated by the fact that conventional ECG electrodes require shaved skin and that awake animals do not easily tolerate skin-attachments unless extensively trained to do so. Hence, ECG in NHPs is either collected from sedated animals [6, 8, 9] or by implanting a telemetry device in combination with highly invasive intracorporal sensors [10–12]. For a non-invasive ECG acquisition from non-sedated monkeys a wearable jacket can be used [1, 13, 14], but this also requires extensive training and physical contact when preparing the animal, which may affect the physiological state.

If heart rate is of interest, but not other ECG parameters, a low-cost alternative to ECG is provided by the photoplethysmogram (PPG). PPG utilizes variations of the light reflected by the skin which correlate with changes of blood volume in the microvascular bed of the skin tissue [15]. PPG allows quite accurate estimation of pulse rate [16–18]. For many applications, pulse rate is a sufficient proxy for heart rate, the latter of which can be, strictly speaking, only assessed from direct cardiac measurements, such as ECG. Conventional PPG still requires a contact sensor comprising a light source to illuminate the skin and a photodetector to measure changes in the reflected light intensity, as used for example in medical-purpose pulse oximeters.

Imaging photoplethysmogram (iPPG) has been proposed [19–21] as a remote and non-contact alternative to the conventional PPG in humans. iPPG is acquired using a video camera instead of a photodetector, under dedicated [19, 22–24] or ambient [21, 25] light. The video is usually recorded from palm or face regions [21–25].

Pilot studies have demonstrated the possibility of extracting iPPG from anesthetized animals, specifically pigs [26, 27]. Since iPPG allows easy and non-invasive estimation of pulse rate, this technique would be very useful for NHP studies. If applicable in non-sedated and behaving animals, it could, for instance, contribute to the welfare of NHP used in research. To our best knowledge, there were no attempts to acquire iPPG from NHPs. In this paper, we demonstrate iPPG extraction from NHP facial videos and provide the first empirical evidence that rhesus monkeys pulse rate can be successfully estimated from iPPG.

## 2 Materials and methods

### 2.1 Animals and animal care

Research with non-human primates represents a small but indispensable component of neuroscience research. The scientists in this study are aware and are committed to the great responsibility they have in ensuring the best possible science with the least possible harm to the animals [28].

Three adult male rhesus monkeys (*Macaca mulatta*) participated in the study (Table 1). All animals had been previously implanted with cranial titanium or plastic “head-posts” under general anesthesia and aseptic conditions, for participating in neurophysiological experiments. The surgical procedures and purpose of these implants were described previously in detail [29, 30]. Animals were extensively trained with positive reinforcement training [31] to climb into and stay seated in a primate chair, and to have their head position stabilized via the head-post implant. This allows implant cleaning, precise recordings of gaze and neurophysiological recordings while the animals work on cognitive tasks in front of computer screen. Here, we made opportunistic use of these situations to record facial videos in parallel. The experimental procedures were approved of by the responsible regional government office (Niedersaechsisches Landesamt fuer Verbraucherschutz und Lebensmittelsicherheit (LAVES)). The animals were pair- or group-housed in facilities of the German Primate Center (DPZ) in accordance with all applicable German and European regulations. The facility provides the animals with an enriched environment (incl. a multitude of toys and wooden structures [32], natural as well as artificial light and access to outdoor space, exceeding the size requirements of European regulations. The DPZ’s veterinarians, the animal facility staff and the lab’s scientists carefully monitor the animals’ psychological and veterinary welfare.

During the video recordings, the animals were head-stabilized and sat in the primate chair in their familiar experimental room, facing the camera. Recording sessions were performed during the preparation phase for the cognitive task. Animals were not rewarded during the video recording to minimize facial motion from sucking, chewing or swallowing.

**Table 1:**
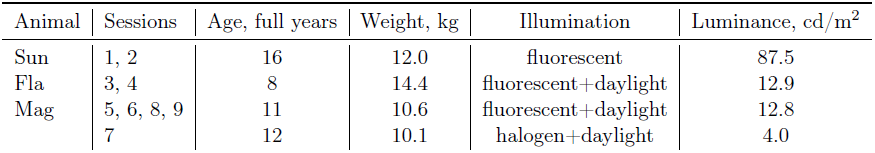
Subjects data and illumination conditions of experimental sessions. For illumination we used either fluorescent lamps mounted at the room ceiling (Philips Master Tl-D 58W/840, 4000K cold white) or setup halogen lamps (Philips Brilliantline 20W 12V 36D 4000K). Luminance was measured by luminance meter LS-100 (Minolta) with close-up lens No 135 aiming at the ridge of the monkey nose (settings: calibration preset; measuring mode abs.; response slow; measurement distance ≈ 55 cm).

| Animal | Sessions | Age, full years | Weight, kg | Illumination | Luminance, cd/m<sup>2</sup> |
| --- | --- | --- | --- | --- | --- |
| Sun | 1, 2 | 16 | 12.0 | fluorescent | 87.5 |
| Fla | 3, 4 | 8 | 14.4 | fluorescent+daylight | 12.9 |
| Mag | 5, 6, 8, 9 | 11 | 10.6 | fluorescent+daylight | 12.8 |
|  | 7 | 12 | 10.1 | halogen+daylight | 4.0 |

### 2.2 Materials and set-up

Our study of pulse rate estimation consisted of nine experimental sessions. Video acquisition parameters for Sessions 1–7 where RGB video was recorded are provided in Table 2. Most methods and results are reported for the RGB video. In addition, monochrome infrared (IR) video was acquired during Session 2 (simultaneously with RGB) and in Sessions 8 and 9, as a control to compare pulse rate estimation from IR and RGB video, see Section 3.5 for details. All videos were acquired at ambient (non-dedicated) light, see Table 1 for details. We used different illumination conditions since setting a particular illumination is often not possible in everyday situations, and we wanted our video-based approach to be able to cope with this variability.

**Table 2:**
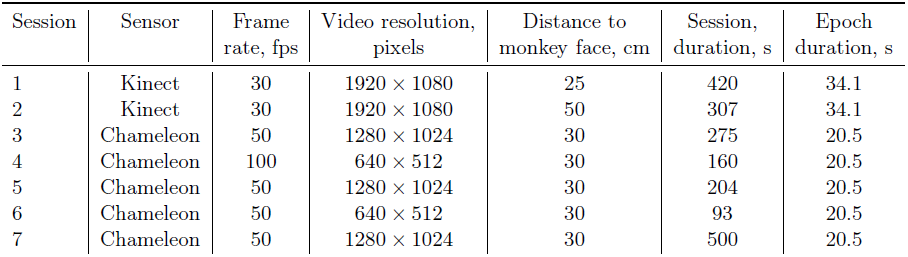
Video acquisition parameters for Sessions 1–7. Kinect here abbreviates the RGB sensor of a Microsoft Kinect for Xbox One; Chameleon stands for Chameleon 3 U3-13Y3C (FLIR Systems, OR, USA) with 10-30mm Varifocal Lens H3Z1014 (Computar, NC, USA) set approximately to 25 mm. Color resolution for both cameras is 8 bit per channel. For Chameleon 3 aperture was fully opened, automatic white-balance was disabled.

| Session | Sensor | Frame rate, fps | Video resolution, pixels | Distance to monkey face, cm | Session, duration, s | Epoch duration, s |
| --- | --- | --- | --- | --- | --- | --- |
| 1 | Kinect | 30 | 1920 × 1080 | 25 | 420 | 34.1 |
| 2 | Kinect | 30 | 1920 × 1080 | 50 | 307 | 34.1 |
| 3 | Chameleon | 50 | 1280 × 1024 | 30 | 275 | 20.5 |
| 4 | Chameleon | 100 | 640 × 512 | 30 | 160 | 20.5 |
| 5 | Chameleon | 50 | 1280 × 1024 | 30 | 204 | 20.5 |
| 6 | Chameleon | 50 | 640 × 512 | 30 | 93 | 20.5 |
| 7 | Chameleon | 50 | 1280 × 1024 | 30 | 500 | 20.5 |

Facial videos of rhesus monkeys were processed to compute iPPG signals. Fig 1 illustrates the main steps of iPPG extraction, which we will consider below in detail: selection of pixels that might contain pulse-related information is described in Section 2.2.1, computation and processing of iPPG is considered in Section 2.2.2.

To estimate pulse rate from iPPG (videoPR) we computed the discrete Fourier transform (DFT) of iPPG signal in sliding overlapping windows; for each window pulse rate is estimated by the frequency with highest amplitude in the heart-rate bandwidth, which is 90–300 BPM (1.5–5 Hz) for rhesus monkeys [2, 33–35]. This approach is commonly used for human iPPG [22, 24, 25], so we applied it here to make our results comparable. The length of DFT window was 1024 points for all sessions except Session 4 where a window length of 2048 points was used due to higher video frame rate. This window length corresponds to 34.1 s for Sessions 1, 2 and 20.5 s for other sessions, see Table 2. A larger window leads to poor temporal resolution, while a smaller window results in crude frequency resolution ( ≥ 4 BPM).

**Figure 1:**
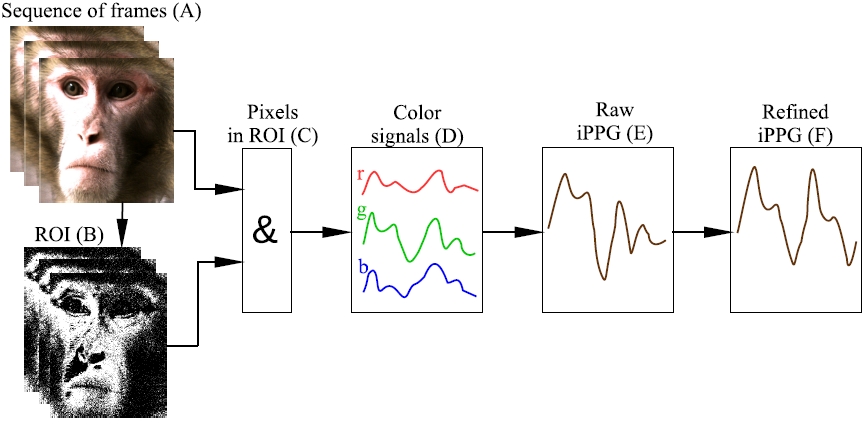
Flowchart of iPPG extraction. (A) We start from a sequence of RGB frames and define for them a region of interest (ROI). (B) For each frame we select ROI pixels containing pulse-related information (shown in white). (C) For these pixels we (D) compute across-pixel averages of unit-free non-calibrated values for red, green and blue color channels. (E) iPPG signal is computed as a combination of three color signals and then (F) refined by using several filters (see main text for detail).

Video processing, ROI selection and computation of color signals are implemented in C++ using OpenCV (opencv.org). iPPG extraction and processing, as well as pulse rate estimation, are implemented in Matlab 2016a (www.mathworks.com).

Reference pulse rate (refPR) values were obtained by a pulse-oximeter CAPNOX (Medlab GmbH, Stutensee) using a probe attached to the monkey’s ear. This pulse-oximeter does not provide a data recording option, but it detects pulses and displays on the screen a value of pulse rate computed as an inverse of the average inter-pulse interval over the last 5 s. The pulse-oximeter measures pulse rate within the range 30–250 BPM with a documented accuracy of ± 3 BPM. The pulse-oximeter screen was recorded simultaneously with the monkeys’ facial video and pulse rate was extracted off-line from the video.

To compare videoPR with refPR, we used epochs of length equal to the length of DFT windows with 50% overlap (see Table 2). For this we averaged the refPR values over the epochs. Note that the quality of the raw iPPG signal was low compared to the contact PPG acquired by the pulse-oximeter. Therefore we were unable to directly detect pulses in the signal, which would provide better temporal resolution (see Section 4.3 for details).

#### 2.2.1 Selecting and refining the region of interest

To reduce spatially uncorrelated noise and enhance the pulsatile signal, we averaged values of color channels over the ROI [21, 36]. In order to select pixels containing maximal amount of pulsatile information, we used the following three-step algorithm:

Step 1. A rectangular boundary for the ROI was selected manually for the first frame of the video. Since no prominent motion was expected from a head-stabilized monkey, this boundary remained the same for the whole video. We tried six heuristic regions shown in Fig 2; the best iPPG extraction for most sessions was achieved for the nose-and-cheeks region 3 (Fig 3).

Step 2. Since hair, eyes, teeth, etc. provide no pulsatile information and deteriorate quality of the acquired iPPG, only skin pixels should constitute the ROI. To distinguish between skin and non-skin pixels, we transformed frames from the RGB to the HSV (Hue-Saturation-Value) color model as recommended in [37,38]. We excluded all pixels having either H, S or V value outside of a specified range. Three HSV ranges describing monkey skin for different illumination conditions (Table 1) were selected by manual adjustment under visual control of the resulting pixel area, see Table 3.

**Figure 2:**
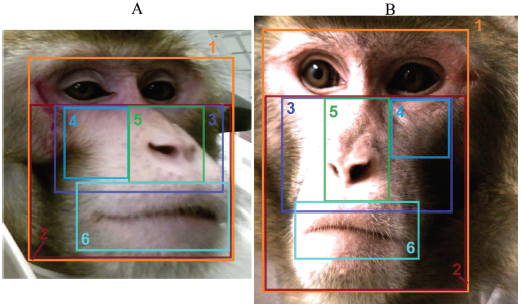
Regions of interest used for iPPG extraction from the video of Sessions 1 (A) and 5 (B): (1) full face, (2) face below eyes, (3) nose and cheeks, (4) nose, (5) cheek, (6) mouth with lips.

**Figure 3:**
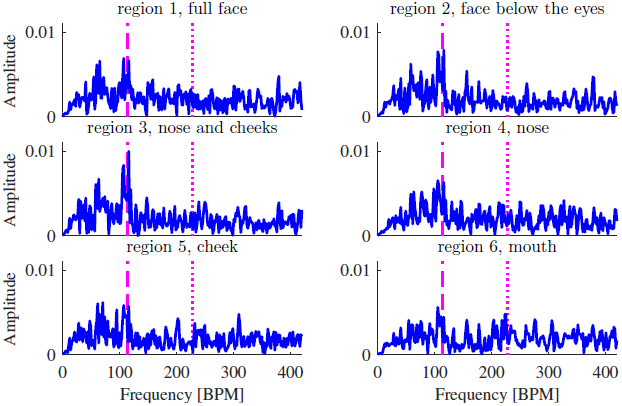
Amplitude spectrum densities of raw iPPG acquired from regions 1-6 (Fig 2) from video of Session 1, 340-374 s. The frequency corresponding to the refPR value and its second harmonic are indicated by dashed and dotted lines, respectively. Although the peak corresponding to the pulse rate is most prominent for region 3, it is distinguishable for other regions as well.

Step 3. In addition, for each frame we excluded all outlier pixels that differed significantly from other pixels in the ROI. The aim of this step was to eliminate pixels corrupted by artifacts. Namely, we excluded pixel (*i*,*j*) in the *k*-th frame if for it the value of any color channel 
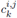
 did not satisfy the inequality

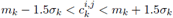

where *m*_*k*_ and σ_k_ are the mean and standard deviation of color channel *c* for pixels included in the ROI of the *k*-th frame at steps 1-2.

**Table 3:**
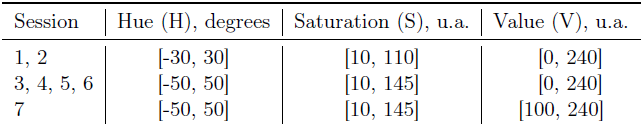
HSV ranges used for skin detection in RGB videos.

#### 2.2.2 Extraction and processing of imaging photoplethysmogram

Color signals 
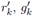
 and 
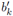
 were computed as averages for each color channel over the ROI for every frame k. Then, prior to iPPG extraction, we mean-centered and scaled each color signal 
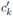
 to make them independent of the light source brightness level and spectrum, which is a standard procedure in iPPG analysis [22,24]:

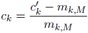

where *m*_*k,M*_ is an M-point running mean of color signal 
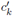

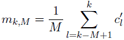

For *k* < *M* we use *m*_1,*M*_. We followed [24] in taking M corresponding to 1 s.

For iPPG extraction we used the “green-red difference” (G-R) method [39], which is simple and effective for computing human iPPG [24, 40]. This method is based on the assumption that green color signal carries maximal amount of pulsatile information, while red color signal contains little relevant information but allows to compensate those artifacts common for both color channels (see Section 4.1 for a discussion of pulsatile information provided by different colors). Thus iPPG was computed as a difference of green and red color signals:

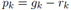

where *g*_*k*_ and *r*_*k*_ are mean-centered and scaled green and red color signals, respectively.

We have also employed two other methods for iPPG extraction, CHROM [22] and POS [24], that are most effective for computing human iPPG [24, 40]. POS computes iPPG as a combination of color signals:

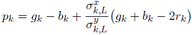

w—here 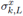 and 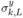 are *L*-point running standard deviations of 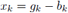 and 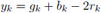 respectively.

CHROM combines color signals in a slightly different way:

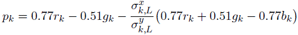

where 
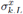
 and 
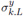

are L-point running standard deviations of 
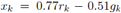
 and 
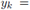

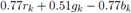
 For both methods we took *L* corresponding to 1 s so that the time window captured at least one cardiac cycle as recommended in [24].

Note that our implementation of POS and CHROM methods is slightly different from the original in [22, 24]: to ensure signal smoothness, we compute running means and standard deviations instead of computing segments of iPPG in several overlapped windows and then gluing them by the overlap-add procedure described in [22]. Processing of iPPG signal for POS and CHROM methods was the same as for G-R method.

Since we used low-cost cameras, we expected a poor quality of the raw iPPG signal compared to contact PPG. Therefore we post-processed iPPG using three following steps, typical for iPPG signal processing [22,25,39,41].

Step 1. We suppressed frequency components outside of the heart-rate bandwidth 90–300 BPM (1.5– 5 Hz) using finite impulse response filter of the 127th order with a Hamming window as suggested in [25].

Step 2. We suppressed outliers in the iPPG signal by cropping amplitude of narrow high peaks that could deteriorate performance on the next step:

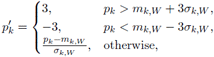

where *m*_*k,W*_ and σ_*k,W*_ are *W*-point running mean and standard deviation of the iPPG signal respectively:

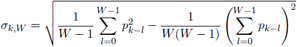

We took *W* equal to 2 s to have estimates of mean and standard deviation over several cardiac cycles.

Step 3. We applied a wavelet filtering proposed in [42] to suppress secondary frequency components. This filtering consists of two steps: first a wide Gaussian window suppresses frequency components remote from the frequency corresponding to maximum of average power over 30 s (global filtering for suppressing side bands that could cause ambiguities in local filtering). Then a narrow Gaussian window is applied for each individual time-point’s maximum (local filtering to emphasize the “true” maximum). As suggested in the original article, we employed Morlet wavelets. We used scaling factors 2 and 5 for global and local filtering, respectively (see [42] for details), since these parameters provide best pulse rate estimation in our case. We implemented wavelet filtering using Matlab functions cwtft/icwtft from Wavelet Toolbox.

To demonstrate the effect of the multi-step iPPG processing, we show in Fig 4 how each processing step magnifies the pulse-related component.

**Figure 4:**
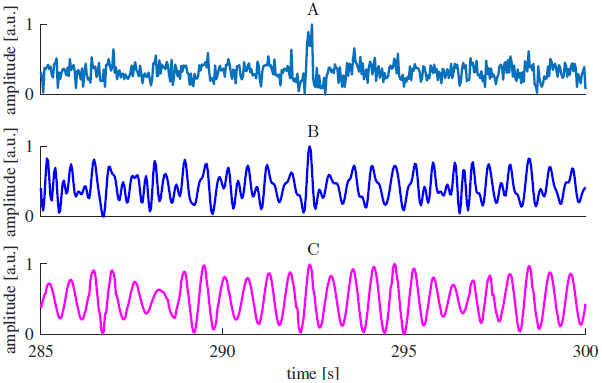
Effect of iPPG processing. (A) Raw iPPG extracted using G-R method from the video of Session 1, (B) iPPG after band-pass filtering, (C) iPPG after wavelet filtering.

## 3 Results

### 3.1 Main result: estimation of pulse rate from RGB video

To check the usefulness of the iPPG signal for pulse rate estimation in rhesus monkeys, we computed several quality metrics characterizing values of pulse rate derived from the video (videoPR) in comparison with the reference pulse rate from the pulse-oximeter (refPR).

The values of quality metrics presented in Table 4 show that pulse rate estimation from iPPG was successful. Bearing in mind that accuracy of refPR is ± 3 BPM, obtained values of mean absolute error for videoPR are rather good. Altogether, for 80% of epochs error of pulse rate estimation was below 7 BPM (which is 5% of rhesus monkey average heart rate 140 BPM), and for 82% of epochs error of pulse rate estimation was below 5% of refPR value. The low correlation between videoPR and refPR for Session 4 is explained by two outlier data-points for which refPR is above average while videoPR is well below average (estimation errors for these epochs are 18 and 11 BPM); after excluding these two points the correlation for Session 4 was 0.82.

**Table 4:** Quality metrics for iPPG-based pulse rate estimates. Error was calculated as the difference between videoPR and refPR; the Pearson correlation coefficient (corr) was computed between videoPR and refPR across epochs in each session.

| Session | Animal | Number of epochs | Mean absolute error, BPM | Epochs with absolute error below |  |  |  | Correlation |
| --- | --- | --- | --- | --- | --- | --- | --- | --- |
|  |  |  |  | 3.5 BPM | 7 BPM | 5% | 10% |  |
| 1 | Sun | 23 | 2.32 | 74% | 100% | 87% | 100% | 0.88 |
| 2 | Sun | 16 | 3.95 | 69% | 75% | 75% | 94% | 0.52 |
| 3 | Fla | 26 | 2.81 | 69% | 96% | 96% | 100% | 0.74 |
| 4 | Fla | 15 | 4.32 | 60% | 80% | 80% | 93% | 0.02 |
| 5 | Mag | 19 | 3.78 | 58% | 89% | 95% | 100% | 0.92 |
| 6 | Mag | 8 | 2.16 | 75% | 100% | 100% | 100% | 0.98 |
| 7 | Mag | 48 | 7.01 | 31% | 56% | 67% | 98% | 0.85 |

Fig 5 shows the Bland-Altman plots for all sessions of each monkey. One can see that for most epochs the error of pulse rate estimation was low; it was slightly higher for high pulse rate.

Fig 6 shows power spectra of the processed iPPG and average values of videoPR for Sessions 1 and 5 in comparison with the reference pulse rate from the pulse-oximeter (refPR). For both sessions, videoPR allows to track the pulse rate changes comparably to the data from pulse-oximeter

### 3.2 Multi-step iPPG processing results in better pulse rate estimation

In this study we used a multi-step procedure for iPPG signal processing. To demonstrate that all the steps are important for the quality of pulse rate estimation, we estimated pulse rate omitting certain processing steps. Average quality metrics for these estimates are presented in Table 5, as one can see only a combination of filters improves the pulse rate estimation.

**Table 5:** Quality metrics averaged over all sessions for different variants of iPPG processing.

| Processing type | Mean absolute error, BPM | Epochs with absolute error below |  | Correlation |
| --- | --- | --- | --- | --- |
|  |  | 3.5 BPM | 7 BPM |  |
| no processing (Fig 4A) | 8.2 | 48% | 72% | 0.48 |
| only band-pass filter (Fig 4B) | 9.9 | 48% | 72% | 0.50 |
| only wavelet filter | 21.4 | 43% | 55% | -0.13 |
| complete processing (Fig 4C) | 4.4 | 56% | 80% | 0.74 |

**Figure 5:**
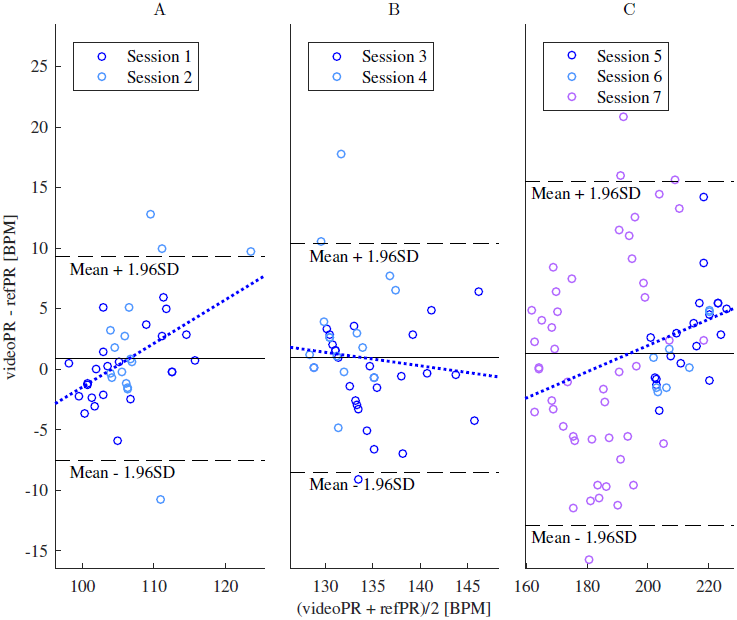
Bland-Altman plots of pulse rate estimates for monkeys Sun (A), Fla (B) and Mag (C). Solid lines show mean error, dashed lines indicate the limits of agreement between videoPR and refPR (mean error ± 1.96 times standard deviation of the error). Dotted lines represent the regression line, regression coefficients were significant for data of Sun (*p* = 0:005) and Mag (*p* = 0:01), indicating that estimation errors slightly increased with pulse rate (this might be partially due to lower precision of refPR for high pulse rate values that are unusual for humans). Regression coefficient for Fla data (B) was insignificant (*p* = 0:13).

**Figure 6:**
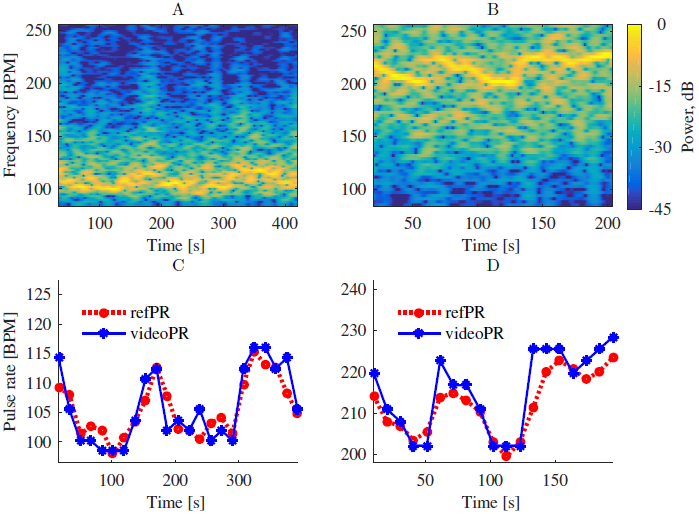
Power spectrum of iPPG (A, B) and pulse rate values (C, D) for Sessions 1 and 5 respectively.

### 3.3 Comparison of methods for iPPG extraction

All results reported so far were obtained using G-R method [39] for iPPG extraction. We have additionally estimated pulse rate from the iPPG signal extracted by CHROM [22] and POS [24] methods. Fig 7 shows that performance of all methods was rather similar, although iPPG extracted by CHROM resulted in a number of large pulse rate estimation errors. Worse performance of CHROM was likely due to the fact that it is based on a model of a human skin [24], thus additional research is required to adjust it for iPPG extraction in NHP.

**Figure 7:**
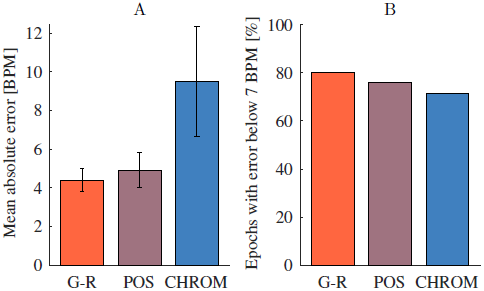
Comparison of iPPG extraction methods. (A) Mean values with 95% confidence intervals (estimated as 1.96 times standard error) of absolute pulse rate estimation error and (B) share of epochs with absolute error below 7 BPM for G-R, POS and CHROM methods across Sessions 1-7.

### 3.4 Effects of motion on iPPG quality

Although we have considered here iPPG extraction for head-stabilized monkeys, our subjects exhibited a considerable number of facial movements. Fig 8 illustrates their negative impact on iPPG extraction quality: the more prominent are movements during an epoch, the lower is quality of extracted iPPG.

**Figure 8:**
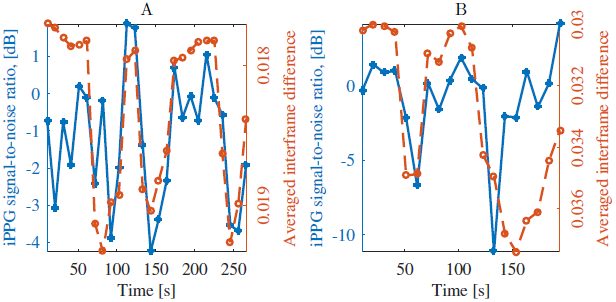
Influence of facial movements on quality of iPPG signal from Sessions 3 (A) and 5 (B). We assess quality of iPPG by signal-to-noise ratio as suggested in [22] (we considered frequency components as contributing to signal if they are in range [refPR − 9 BPM, refPR + 9 BPM] or [2refPR − 18 BPM, 2refPR + 18 BPM]). We estimate amount of motion by averaging mean squared differences between successive frames. In epochs with more motion, iPPG signal-to-noise ratio is mostly lower. (Note the inversed y-axis for motion.)

### 3.5 Estimation of pulse rate from infrared video

To check the possibility of iPPG extraction from monochrome infrared (IR) video in NHP, we recorded video using the Kinect IR sensor during Session 2. Additionally, we conducted Sessions 8 and 9 (see Table 1) using the monochrome camera Chameleon 3 U3-13Y3M (FLIR Systems, OR, USA) with ultraviolet/visual cut-off filter R-72 (Edmund Optics, NJ, USA) blocking light with wavelengths below 680 nm. Details of the recorded video are presented in Table 6. Extraction of iPPG from monochrome IR video was similar to that from RGB video: we computed raw iPPG as average intensity over all pixels in a ROI (monochrome IR video provides a single intensity channel, so we could not use G-R method in this case) and then processed the signal as described in Section 2.2.2. For ROI selection we determined skin pixels as having intensity within the range specified in Table 6.

**Table 6:** Description of experimental sessions with monochrome infrared video recording.

| Session | Sensor | Frame rate, fps | Video resolution, pixels | Distance to monkey face, cm | Session, duration, s | Skin filter |
| --- | --- | --- | --- | --- | --- | --- |
| 2 | Kinect IR | 30 | $512 \times 424$ | 50 | 307 | 100–253 |
| 8 | Chameleon | 100 | $1280 \times 1024$ | 30 | 133 | 80–240 |
| 9 | Chameleon | 50 | $1280 \times 1024$ | 30 | 184 | 80–240 |

As indicated by the quality metrics in Table 7, pulse rate estimation from IR video was not successful. The low quality of estimation reflects that pulse rate had most of the time lower power than other frequencies related to movements, lighting variations, other artifacts or noise.

**Table 7:** Quality metrics for pulse rate estimates computed from monochrome IR video.

| Session | Number of epochs | Mean absolute error, BPM | Epochs with absolute error below |  | Correlation |
| --- | --- | --- | --- | --- | --- |
|  |  |  | 3.5 BPM | 7 BPM |  |
| 2 | 16 | 52.5 | 13% | 19% | -0.02 |
| 8 | 12 | 37.4 | 25% | 42% | -0.13 |
| 9 | 17 | 97.3 | 6% | 6% | -0.01 |

## 4 Discussion

### 4.1 Imaging photoplethysmography under visible and infrared light

Quality of the photoplethysmogram (PPG) acquisition in general strongly depends on the light wavelength. The PPG signal is generated by variations of the reflected light intensity due to changes in scattering and absorption of skin tissue. In the ideal case, light would be only absorbed by blood haemoglobin, which would make PPG a perfect indicator of the blood volume changes [16,43]. However, light is also absorbed and scattered by water, melanin, skin pigments and other substances [16].

There is no general consensus about the optimal wavelength for PPG acquisition, but the trend is in favor of employing visible light [18, 44]. Although traditionally PPG was acquired at near infrared (NIR) wavelengths [15, 16], experiments have shown that the light from the visible spectral range allows to acquire comparably accurate PPG [43, 45, 46] or to even reach higher accuracy [47, 48]. Theoretical studies and simulations in [39] indicate that the best signal-to-noise ratio in iPPG should be obtained for wavelengths in the ranges of 420–455 and 525–585 nm, with peaks at 430, 540 and 580 nm corresponding to violet, green and yellow light, respectively (see [39, Section 3.5] for details). Passing through epidermis and bloodless skin is optimal for green (510–570 nm), red (710–770 nm) and NIR light (770–1400 nm) [49, 50]. However, absorption of haemoglobin is maximal for the green and yellow (570–590 nm) light [47, 51, 52]. This results in a better signal-to-noise ratio for these wavelengths [18,44,47].

Our results show that monochrome infrared (IR) video acquired with single non-specialized cameras and without dedicated illumination was not suitable for iPPG extraction, while RGB visible-light video was suitable. The striking difference between the results is not altogether surprising. Combining several color channels when extracting iPPG from RGB video, is more effective than considering data from a single wavelength [21,22]. Specifically, it makes iPPG extraction from RGB video less sensitive to movements and lighting variations, while for IR video to compensate them simultaneous video acquisition from several cameras is typically used [53,54].

Additional research is however required to check whether visible or IR light is generally preferable for iPPG extraction in NHP. In particular, two following obstacles may hinder accurate iPPG extraction from RGB video of NHP faces under visible light:

- **High melanin concentration.** Melanin strongly absorbs visible light with wavelengths below 600 nm [16], which degrades the quality of PPG for humans with a high melanin concentration [41, 55]. This often motivated using red or NIR light for PPG acquisition [16, 17], but some modern methods for iPPG extraction successfully alleviate this problem [55]. To the best of our knowledge, melanin concentration in monkey facial skin was not compared to that of humans, and the question of melanin concentration influence on iPPG extraction in NHP remains open.
- **Insufficient light penetration depth.** For a light of wavelength λ within 380–950 nm (visible and NIR light) with fixed intensity, the higher λ is, the deeper the light penetrates the tissue [46, 56]. This is important since pulsatile variations are more prominent in deeper lying blood vessels. For instance, the penetration depth of the blue light (400-495 nm) is only sufficient to reach the human capillary level [39, 57], providing little pulsatile information [23]. Green light penetrates up to 1 mm below the human skin surface, which makes the reflected light sensitive to the changes in the upper blood net plexus and in the reticular dermis [39,56]. To assess blood flow in the deep blood net plexus, one uses red or NIR light with penetration depth above 2 mm [21, 56]. Insufficient penetration depth of visible light may hinder iPPG acquisition from certain parts of human body [19, 39], but human faces allow extraction of accurate iPPG from the green and even the blue light component [21,46].

For rhesus monkeys iPPG extraction is simplified by the fact that their facial skin has a dense superficial plexus of arteriolar capillaries [58]. It has been previously demonstrated [59] that subtle changes in facial color of rhesus monkeys caused by blood flow variations can be detected using a camera sensor. For other NHP it is not clear which wavelength if any would be sufficient to penetrate the skin and systematic studies are required to answer this question.

### 4.2 Video acquisition and region of interest selection

A good video sensor is necessary for successful iPPG extraction. In this study we used three different sensors: RGB and IR sensors of a Microsoft Kinect for Xbox One and Chameleon 3 1.3 MP (color and monochrome models), details are provided in Table 8.

**Table 8:** Characteristics of sensors used for the experiments (according to [60–62]).

| Camera | Max. video resolution, pixels | Pixel size, $\mu m$ | Max. frame rate, fps |
| --- | --- | --- | --- |
| Kinect RGB | 1920x1080 | 3.1 | 30 |
| Kinect IR | 512x424 | 10 | 30 |
| Chameleon 3 1.3 MP | 1280x1024 | 4.8 | 149 |

Although all these CMOS-sensors have moderate characteristics, they are better than sensors of most general-purpose cameras used for iPPG extraction in humans (see, for instance, [25, 63]). Still, specialized cameras, like those used in [23,39], could provide more robust and precise iPPG extraction in NHP. For the choice of the camera such characteristics as pixel noise level and sensitivity are crucial, since iPPG extraction implies detection of minor color changes. Besides, sufficient frame rate is required to capture heart cycles. For humans the minimal frame rate for iPPG extraction is 15 fps [25], though in most cases 30 fps and higher are used [63]. Since heart rate for rhesus monkeys is almost twice as high as for humans, higher frame rate is required. In our study frame rate of 30 fps (provided by Microsoft Kinect) was sufficient to estimate pulse rate from iPPG; in general we expect that 50 fps should be enough for accurate iPPG extraction in rhesus monkeys.

For practical applications, one needs a criterion for rejecting video frames if they contain too little reliable pulsatile information for accurate iPPG extraction. One possible solution is to reject video frames as invalid if the signal-to-noise ratio of the acquired iPPG is not above a threshold computed from ROIs that do not contain exposed skin (e.g. regions covered by hair). See also [64] for a method of rejecting invalid video based on spectral analysis.

In this paper we have only considered manual selection of ROI, which is acceptable for head-stabilized NHPs. In the general case, automatic face tracking would be of interest [65]. In this study, a cascade classifier using the Viola-Jones algorithm [66] allowed detection of monkey faces in recorded video. Furthermore, techniques for automatic selection of regions providing most pulsatile information in humans [36, 67] can be adopted.

For non-head-stabilized NHP, robustness of the iPPG extraction algorithm to motion is especially important. For humans this problem can be successfully solved by separating pulse-related variations from image changes reflecting the motion [68], but for NHP additional research is required. In particular, hair covering parts of NHP face can provide references for successful compensation of small motion (see [39], where similar methods are proposed for humans).

An even more challenging task is iPPG extraction for freely moving NHP. In this case distance of the animal face to camera sensor and relative angles to the sensor and to the light sources are time dependent. For humans, a model taking this into account was suggested in [24], however its stable performance has been only demonstrated for relatively short (several meters) and nearly constant distances from the face to the camera. Nevertheless, recent progress in iPPG extraction from a long distance video [63] and in compensation of the variable illumination angle and intensity [24, 69] gives hope that required models might be developed in future.

### 4.3 Pulse rate estimation

In our study estimated pulse rates were in the range of 95–125 BPM for monkey Sun, 125–150 BPM for Fla and 160–230 BPM for Mag, which agrees with previously reported values of heart rate (120-250 BPM) for rhesus monkeys sitting in a primate chair [2, 33–35]. Performance of our method for pulse rate estimation (Table 4) was only slightly worse than those reported for humans: mean absolute error obtained in [70] was 2.5 BPM; fraction of epochs with error below 6 BPM (about 8% of average human pulse rate) for the best method considered in [71] was achieved for 87% of epochs; reported values of the Pearson correlation coefficient vary from 0.87 in [71] to 1.00 in [25]. Comparing our results with outcomes of human studies, one should also allow for the imperfect reference since data from pulse-oximeter are not as reliable as ECG.

In this study we used the discrete Fourier transform (DFT) for pulse rate estimation. Despite of its popularity this method is often criticized as being imprecise [19, 71]). Indeed, application of DFT implies analysis of iPPG signal in a rather long time window (20-40 s), while pulse rate is non-stationary and scarcely remains constant for several heart beats in row [19]. These fluctuations blur the iPPG spectrum and may hinder pulse rate estimation.

As an alternative to using DFT, we estimated instantaneous pulse rate from inter-beat intervals defined either as difference between adjacent systolic peaks (maximums of iPPG signal) or as diastolic minima (troughs of the signal) [7]. Since the wavelet filter greatly smoothed the signal and modified its shape (see Fig 4), we excluded this processing step. We tested several methods for peak detection in PPG [72, 73], but the results were inconclusive and highly dependent on the quality of iPPG signal, therefore we do not present them here. Further research is required to find suitable algorithms for estimation of inter-beat intervals from iPPG in NHP. Specifically, the choice of methods for iPPG signal processing prior to peak detection seems to be important. Note that the post-processing applied here destroys the shape of pulse waves (see Fig 4). However, in this study the quality of iPPG signal was only sufficient for a rough frequency-domain analysis, thus the shape of iPPG signal was not critical.

### 4.4 Summary

Here we evaluated and documented the feasibility of imaging photoplethysmogram (iPPG) extraction for pulse rate estimation in rhesus monkeys. Our future work will focus on enhancement of iPPG extraction quality, development of a method for robust iPPG extraction in NHP without head-stabilization and investigating the possibility of effective pulse rate estimation from infrared video.

In summary, we demonstrated the possibility of imaging photoplethysmogram extraction from RGB facial video of head-stabilized macaque monkeys. For the video recording we used general-purpose cameras and ambient artificial light. Our results show that one can estimate, accurately and non-invasively, the pulse rate of awake NHPs without special hardware for illumination and video acquisition and using only standard algorithms of signal processing. This makes imaging photoplethysmography a promising tool for non-invasive and non-contact remote estimation of pulse rate in non-human primates, applicable for behavioral and physiological studies of emotion and social cognition, as well as for the welfare-assessment of animals in research.

## Acknowledgments

The authors would like to thank Dr. D. Pfefferle for the help with data acquisition.

We acknowledge funding from the Ministry for Science and Education of Lower Saxony and the Volkswagen Foundation through the program “Niedersächsisches Vorab” and the DFG research unit 2591 “Severity assessment in animal based research”. Additional support was provided by the Leibniz Association through funding for the Leibniz ScienceCampus Primate Cognition and the Max Planck Society.

